# Prenatal exposure to environmental stressors alters gut macrophage development and gastrointestinal function of male offspring

**DOI:** 10.1101/2025.03.24.644910

**Authors:** Dang M. Nguyen, Sarah K. Monroe, Danielle N. Rendina, Kevin S. Boyd, Erika D. Rispoli, Olivia M. Wirfel, A. Brayan Campos-Salazar, Anna R. Araujo, Trisha V. Vaidyanathan, Virginia L. Keziah, Benjamin A. Devlin, Caroline J. Smith, Staci D. Bilbo

**Affiliations:** Department of Psychology and Neuroscience, Duke University, Durham, NC, USA; Department of Neurobiology, Duke University Medical Center, Durham, NC, USA; Howard Hughes Medical Institute; Department of Cell Biology, Duke University Medical Center, Durham, NC, USA

**Keywords:** air pollution, maternal stress, small intestine, gut macrophages, enteric nervous system, neurodevelopmental disorders, gastrointestinal dysfunction, gut-brain axis

## Abstract

Gastrointestinal (GI) dysfunction is a frequently reported comorbidity of neurodevelopmental disorders (NDDs). Early-life inflammatory challenges from the environment (e.g. infection, toxicants) can increase risk for NDDs but the impact of such stressors on the developing GI tract is not well understood. We investigated possible mechanisms by which GI comorbidities occur in response to environmental stressors using our well-characterized model of combined gestational exposure to air pollution (diesel exhaust particles, DEP) and maternal stress (MS), which induces social deficits in male but not female offspring. We show that DEP/MS disrupts normal GI development, leading to altered small intestine morphology in neonatal males, but not females. Recent evidence shows that resident macrophages of the gut prune enteric neurons during a precise postnatal window. We found decreased pruning of gut enteric neurons by the resident macrophages of the muscularis externa in DEP/MS exposed males at postnatal day 14. In line with this, we saw the expression of motor neuron-associated genes spike in males at the same postnatal time point following DEP/MS exposure. Finally, we assessed the motor function of the GI tract of these animals and observed dysmotility in DEP/MS males only. Taken together, these findings establish intestinal macrophages as a mediator of GI development that is sensitive to early-life perturbations from the environment, highlighting a potential mechanism connecting NDDs with comorbid GI dysfunction.

## Introduction

Environmental factors are critical determinants of developmental outcomes. Growing evidence indicates that gestational exposure to immune challenges—such as infections, toxicants, or maternal health complications—increases an individual’s susceptibility to neurodevelopmental disorders (NDDs) such as autism^1–3^. Importantly, recent evidence has emerged that many of the underlying mechanisms by which diverse environmental factors impact neural development involve changes within the gut-brain axis ^4–6^.

Gastrointestinal (GI) abnormalities - e.g. gastritis, diarrhea, acid reflux, etc. – make up some of the most common comorbidities in NDDs, with the majority of diagnosed individuals reporting at least one GI symptom^7,8^. Symptoms of GI dysfunction often coincide with behavioral symptom onset and severity in NDDs^9^, suggesting a coupling between brain development and GI development. However, the biological basis of this connection remains unclear, particularly in the context of environmental exposures.

We have recently demonstrated that male mice exposed prenatally to a combination of air pollution and maternal stress—two of the most consistently highlighted risk factors for NDDs in global health studies^10,11^—exhibit social deficits consistent with many NDDs, whereas females are resilient to these exposures. The male-specific behavioral deficits in this model are driven by impaired microglial pruning of developing thalamocortical synapses during the early postnatal period^12^. The social deficits observed in males were accompanied by significant changes to gut architecture and microbiome, and normalization of the gut microbiome at birth via cross-fostering prevented the social deficits later in life^6^. These results, along with a burgeoning literature examining the gut microbiome and vagal connections to the gut ^4,5,13–15^, suggest that the GI tract is a critical modulator of neuroimmune perturbations and an attractive therapeutic target for NDDs. However, little is known about the cellular mechanisms underlying GI comorbidities in NDDs.

Intestinal macrophages impact GI development via diverse functions including the regulation of barrier integrity through release of inflammatory signals^16^, sculpting gut architecture via apoptotic clearance^17^, and newly discovered interactions with enteric neurons ^18,19^. Specifically, resident macrophages of the muscularis externa prune enteric neuron synapses precisely at postnatal day (P)14, a function analogous to that of microglia within the postnatal brain. While microglial synapse pruning has been demonstrated to have a critical role in brain development and behavioral outcomes, the functional consequences of enteric synapse pruning have not been reported. Notably, some intestinal macrophages in the first postnatal weeks of life have an embryonic yolk sac origin in common with microglia^20^. These data suggest the intriguing possibility that gestational immune (re)programming of these long-lived resident macrophage populations in response to environmental factors could have a shared impact on the developing brain and gut.

In this study, we examined the effects of combined prenatal stressors on intestinal macrophages and GI function within the mouse small intestine during the early postnatal period (P4 and P14). We identify male-specific responses to combined prenatal exposures by intestinal macrophages, along with transcriptional changes associated with gut motor function starting at the second postnatal week of life. We also found a striking male-specific deficit in macrophage refinement of synapses within the muscularis externa, which is predominantly innervated by motor neurons coordinating smooth muscle contractions. Finally, in prenatally stressed males at P14 we observed accelerated GI transit, a symptom of enteric neuron hyperactivity^21^. Together these findings elucidate potential mechanisms by which GI dysmotility can be induced by environmental stressors.

## Results

### Prenatal exposure to combined stressors alters male small intestine morphology, but not permeability

To model pervasive environmental stressors, we used combined gestational exposure to air pollution and stress. We exposed pregnant C57BL6/J dams to intermittent diesel exhaust particle (DEP) tracheal instillations to mimic chronic air pollution as we have reported previously^12^. DEP is a major toxic component of air pollution and a potent inflammatory stimulus^22–24^. We additionally restricted the bedding and nesting material during the last trimester of pregnancy, a previously defined paradigm that induces maternal stress (MS) (i.e., DEP/MS condition) (**Fig 1a**)^25,26^. Control (CON) dams received instillations of the vehicle solution (PBST) and were housed in standard cages with full nesting material. We have previously reported that combined DEP and MS, but neither exposure in isolation, induces a sterile maternal immune response and significant changes in offspring gut and brain function, primarily in males^6,12^.

**Figure 1.**
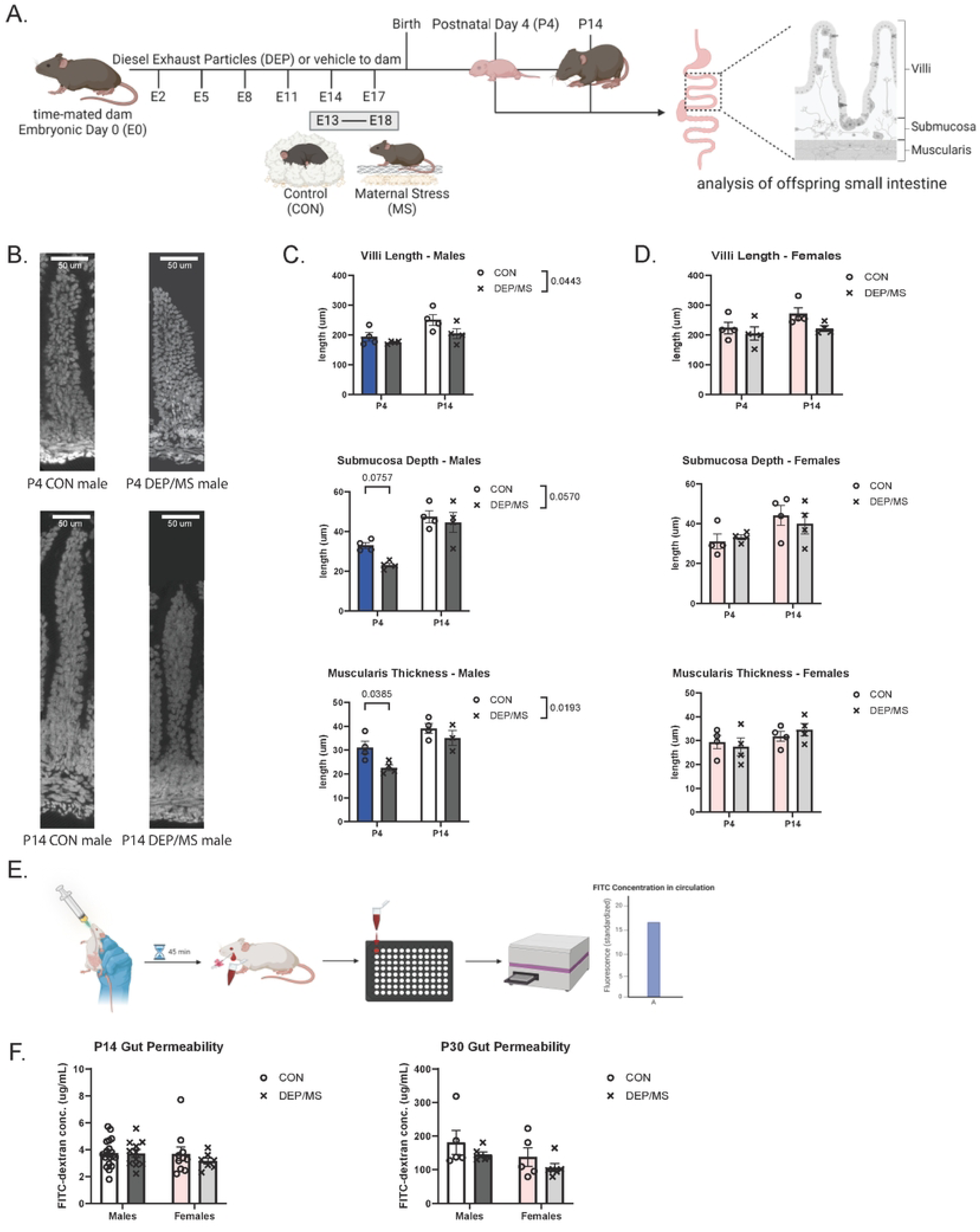
DEP/MS alters small intestine morphology, but not permeability in male offspring. (A) Schematic of DEP/MS paradigm. Time mated dams experienced repeated diesel exhaust particle exposure throughout pregnancy combined with limited bedding and nesting, or control conditions. Offspring tissues were collected at P4 or P14 for analysis (B) Representative images of gut villi in the ileum of male offspring at P4 and P14 using a DAPI stain. (C) Quantification of tissue architecture size in P4 and P14 male offspring in villi (n = 4 mice/group, 2-way ANOVA [treatment x age], treatment: p = 0.0443, age: p = 0.0109, interaction: p = 0.3615), submucosa (n = 4 mice/group, 2-way ANOVA [treatment x age], treatment: p = 0.0570, age: p < 0.0001, interaction: p = 0.2613), and muscularis (n = 4 mice/group, 2-way ANOVA [treatment x age], treatment: p = 0.0193, age: p = 0.0009, interaction: p = 0.3498) (D) Quantification of tissue architecture size in P4 and P14 female offspring in villi (n = 4 mice/group, 2-way ANOVA [treatment x age], treatment: p = 0.0931, age: p = 0.1106, interaction: p = 0.4225), submucosa (n = 4 mice/group, 2-way ANOVA [treatment x age], treatment: p = 0.8043, age: p = 0.0315, interaction: p = 0.4617), and muscularis (n = 4 mice/group, 2-way ANOVA [treatment x age], treatment: p = 0.8931, age: p = 0.1196, interaction: p = 0.4171) (E) Schematic of gut permeability assay. Offspring received FITC-dextran by oral gavage at P14 and P30. After 45 minutes, serum was collected and measured on a microplate for fluorescent intensity of FITC-dextran (F) Quantification of gut permeability assay at P14 (n = 7-21 mice/group, 2-way ANOVA [treatment x sex], treatment: p = 0.4570, sex: p = 0.3767, interaction: p = 0.4316) and P30 (n = 5 mice/group, 2-way ANOVA [treatment x sex], treatment: p = 0.1445, sex: p = 0.0767, interaction: p = 0.9098)

The architecture of the gut wall determines much of the functionality of the GI tract, such as the amount of surface area available for digestion and nutrient absorption. We have previously found DEP/MS offspring have longer villi^6^, mucosal protrusions which interface directly with the small intestine luminal contents, at P45. To see if this difference in villi arises early in development, we measured the architecture of the offspring small intestine at both P4 and P14.

Beneath the villi lie the submucosal and muscularis externa layers of the gut wall – the former of which contains a mix of connective tissue, lymphoid tissue, vasculature, and diverse neuronal ganglia, while the latter consists primarily of smooth muscle tissue and the ganglia of neurons innervating it. We observed DEP/MS male offspring to have significantly shorter villi and a thinner muscularis, along with a trending decrease in submucosal depth, than CON males (**Fig 1b-c**). These male-specific deficits were more pronounced at P4, and we saw no differences in the small intestine morphology of female offspring (**Fig 1d**). We did not see any coinciding effects on the body weight of P4 DEP/MS offspring that could underlie the architectural findings (Fig S1a). At P14, we observed both male and female DEP/MS offspring to have heavier body weights, rather than lighter, without any differences to male small intestine weight and length post-normalization (Fig S1b-g). These results indicate that prenatal DEP/MS induces male-specific impacts on GI morphology early in development.

We next tested for differences in the barrier integrity of the small intestine early in life based on common reports of a “leaky gut” phenotype in NDDs^27,28^. We administered a fluorescent FITC-dextran solution orally to P14 and P30 mice, and after 45 minutes, we measured the amount of FITC-dextran that translocated across the intestinal epithelium into circulation (**Fig 1e**). We did not see any changes in circulating FITC-dextran levels between DEP/MS and CON offspring (**Fig 1f**), suggesting that DEP/MS does not impact permeability of the GI tract during early development. Furthermore, there were no significant differences in the expression of tight junction and inflammatory cytokine genes that modulate intestinal permeability^4,27,28^ (**Fig S2**), demonstrating that morphological changes in response to DEP/MS do not correspond with overt disruptions to epithelial function nor active inflammation in the small intestine.

### DEP/MS impacts male intestinal macrophage surface antigen expression, but not population size

Having observed architectural changes in the small intestine of DEP/MS males, we next assessed GI differences at the cellular level. In the small intestine, the sensitivity of intestinal macrophages following early-life insults remains largely unknown, obscured by diverse subpopulation variability that is enforced by layer-specific local cues. We assessed these different cell populations by measuring their expression of F4/80—a macrophage surface receptor whose homeostatic expression can vary due to prior immune activation ^30,31^, —within each layer of the small intestine at P4 and P14 (**Fig 2a**). Once again, we found male-specific effects of DEP/MS, where F4/80 expression was diminished across both P4 and P14 in comparison to CON males (**Fig 2b-c**), while female intestinal macrophages displayed no differences in F4/80 expression (**Fig 2d**). We also quantified the number of F4/80+ cells in the different layers of the gut wall. We found no significant differences in macrophage density following DEP/MS exposure across all layers of the female small intestine, with similar observations in the male submucosa and muscularis (Fig S3a-b). Within the male villi, DEP/MS only induced a slight increase in macrophage density at P4 that reversed by P14 (Fig S3a). Together, these data suggest that while macrophage number is relatively stable in the male small intestine, the responsivity or function of these cells may be altered by DEP/MS exposure.

**Figure 2.**
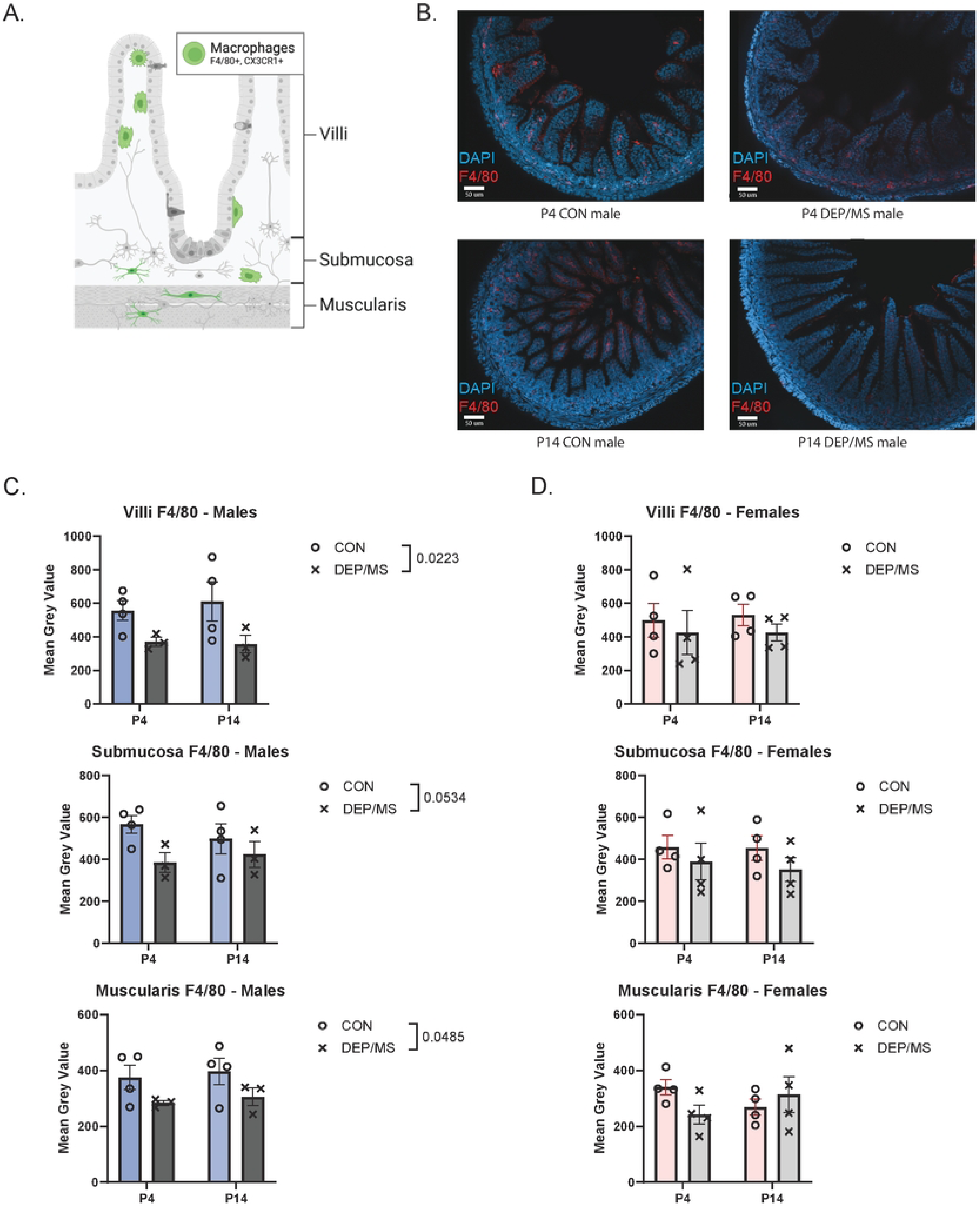
DEP/MS reduces expression of F4/80 expression across intestinal layers in male offspring. (A) Schematic of macrophage distribution throughout intestinal tissue layers. (B) Representative images of F4/80 signal in macrophages of male offspring at P4 and P14. (C) Quantification of F4/80 signal in P4 and P14 male offspring in villi (n = 3-4 mice/group, 2-way ANOVA (treatment x age), treatment: p = 0.0223, age: p = 0.8105, interaction: p = 0.6880), submucosa (n = 3-4 mice/group, 2-way ANOVA [treatment x age], treatment: p = 0.0534, age: p = 0.8042, interaction: p = 0.3821), and muscularis (n = 3-4 mice/group, 2-way ANOVA [treatment x age], treatment: p = 0.0485, age: p = 0.6017, interaction: p = 0.9886) (D) Quantification of F4/80 signal in P4 and P14 female offspring in villi (n = 4 mice/group, 2-way ANOVA [treatment x age), treatment: p = 0.3492, age: p = 0.8693, interaction: p = 0.8663), submucosa (n = 4 mice/group, 2-way ANOVA (treatment x age), treatment: p = 0.2230, age: p = 0.7518, interaction: p = 0.8022), and muscularis (n = 4 mice/group, 2-way ANOVA [treatment x age], treatment: p = 0.5304, age: p = 0.9891, interaction: p = 0.1126)

### DEP/MS is followed by increased motor neuron gene expression specifically at P14 in males

Based on our findings that DEP/MS blunts F4/80 expression in the male small intestine, and our previous work^6,12^ showing coincident neuroimmune deficits in the brain, we next examined the possibility that the enteric nervous system is also altered by DEP/MS exposure. To test this, we first measured neuronal gene expression at P4 and P14 in the male small intestine.

Like the brain, the enteric nervous system has a tight spatial organization, with the majority of its fibers and neuronal cell bodies collecting into two dense plexuses located in the submucosa and muscularis respectively^32^. Over half of these neurons are motor neurons that coordinate contractility of the GI tract and sequester primarily in the muscularis, containing the muscle-flanked myenteric plexus^33^. The other major neuronal subtype is primary afferents, or sensory neurons, which compromise around a quarter of enteric neurons and are represented in both the myenteric and submucosal plexuses (**Fig 3a**). To capture holistic changes to the enteric nervous system, we first looked at the expression of the pan-neuronal synaptosomal-associated protein 25 gene (*SNAP25*), which facilitates fusion of synaptic vesicles to the presynaptic membrane during neurotransmitter release. While DEP/MS exposure had no effect on *SNAP25* expression in the small intestine of P4 males, there was a striking upregulation of *SNAP25* at P14 in the DEP/MS males suggesting increased presynaptic terminal activity emerging by the third week of life (**Fig 3b**).

**Figure 3.**
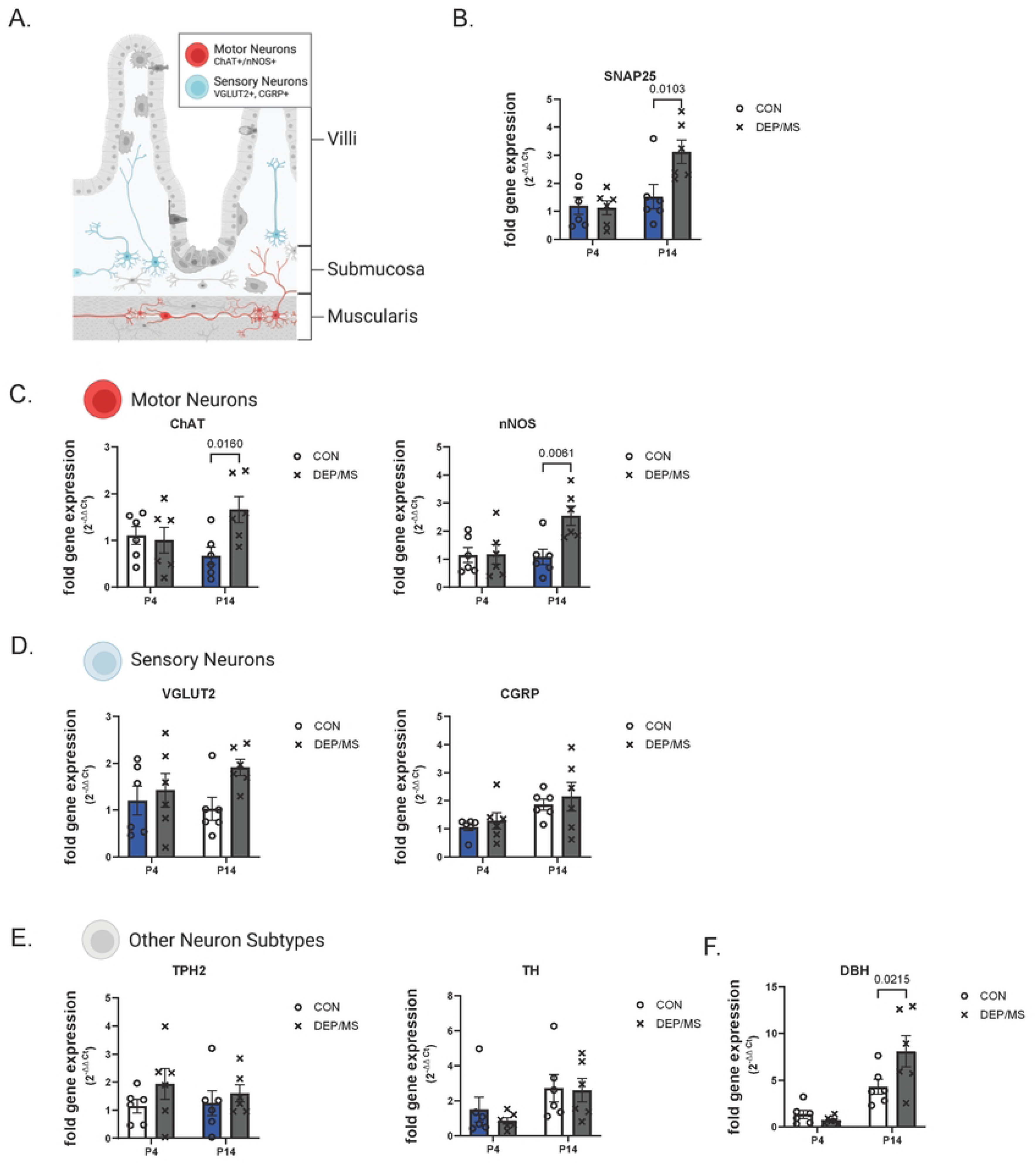
DEP/MS increases expression of motility-related genes in male offspring. (A) Schematic of neuronal subpopulation primary distribution throughout intestinal tissue layers. (B) Quantification of pan-neuronal *SNAP25* gene expression in the small intestine of male offspring (n = 6 mice/group, 2-way ANOVA [treatment x age], treatment: p = 0.0469, age: p = 0.0043, interaction: p = 0.0311) (C) Quantification of motor neuron *ChAT* gene expression (n = 6 mice/group, 2-way ANOVA (treatment x age), treatment: p = 0.0735, age: p = 0.6510, interaction: p = 0.0331) and *nNOS* gene expression (n = 6 mice/group, 2-way ANOVA (treatment x age), treatment: p = 0.0249, age: p = 0.0460, interaction: p = 0.0296) in the small intestine of male offspring (D) Quantification of sensory neuron *VGLUT2* gene expression (n = 6 mice/group, 2-way ANOVA [treatment x age], treatment: p = 0.0616, age: p = 0.5855, interaction: p = 0.2443) and *CGRP* gene expression (n = 6 mice/group, 2-way ANOVA [treatment x age), treatment: p = 0.4288, age: p = 0.0168, interaction: p = 0.9191) in the small intestine of male offspring (E) Quantification of serotonergic neuron *TPH2* gene expression (n = 6 mice/group, 2-way ANOVA [treatment x age), treatment: p = 0.1764, age: p = 0.7785, interaction: p = 0.5848) and dopaminergic neuron *TH* gene expression (n = 6 mice/group, 2-way ANOVA [treatment x age], treatment: p = 0.5536, age: p = 0.0281, interaction: p = 0.6842) in the small intestine of male offspring (F) Quantification of smooth muscle contractility modulator *DBH* gene expression in the small intestine of male offspring (n = 6 mice/group, 2-way ANOVA [treatment x age), treatment: p = 0.1112, age: p < 0.0001, interaction: p = 0.0319)

To assess the specificity of this effect, we analyzed genes associated with the different neuronal subtypes. Choline acetyltransferase (*ChAT*) and neuronal nitric oxide synthase (*nNOS*) are genes involved in the synthesis of acetylcholine and nitric oxide respectively, capturing the excitatory and inhibitory motor neuron populations in the small intestine, along with sensory afferents in the case of *ChAT*^33^. In line with our SNAP25 data, both *ChAT* and *nNOS* expression in the male small intestine were unchanged by DEP/MS at P4, but increased significantly in response to DEP/MS at P14 (**Fig 3c**). This suggests dysregulation of a motor neuron developmental window in response to DEP/MS. We also looked at the expression of vesicular glutamate transporter 2 (*VGLUT2*) and calcitonin gene-related peptide (*CGRP*), which have both been shown to label sensory neurons in the small intestine^34,35^. Neither gene showed significant changes in expression at P4 or P14 following DEP/MS exposure (**Fig 3d**).

Similarly, looking at serotonergic populations via tryptophan hydroxylase 2 (*TPH2*) and dopaminergic populations via tyrosine hydroxylase (*TH*), we saw no effect of DEP/MS on these minor neuronal subtypes in the small intestine (**Fig 3e**). However, dopamine beta-hydroxylase (*DBH*) expression mirrored that of motor neuron genes, with a sharp increase at P14 in DEP/MS males compared to P14 CON males (**Fig 3f**). DBH from the small intestine is utilized in the synthesis of norepinephrine by postganglionic sympathetic nerves, which are extrinsic to the GI tract yet also modulate smooth muscle contractility^36^. Given that DEP/MS effects on the male enteric nervous system converge on motor control, we measured the gene expression of muscle-specific kinase (*MuSK*), a postsynaptic component of the neuromuscular junction, to check if smooth muscles cells were also impacted by DEP/MS exposure (Fig S4a). We did not find DEP/MS to alter *MuSK* expression at P4 or P14, suggesting that motor control differences observed at P14 are specific to the neurons innervating the muscularis (Fig S4b).

### DEP/MS impairs intestinal macrophage-mediated synaptic engulfment within the male muscularis

To identify possible mechanisms responsible for the neuronal perturbations observed in male small intestine following DEP/MS exposure, we investigated pathways by which macrophages and neurons are known to interact in the gut. The most defined function of intestinal macrophages is the release of inflammatory cytokines to combat pathogenic threats; this comes at the expense of triggering apoptosis in neighboring cells including neurons^19^. In line with our observations that DEP/MS reduced macrophage reactivity rather than increasing it, we did not observe differences in pro-inflammatory interleukin-1 beta (*IL-1β*), interleukin-6 (*IL-6*), nor tumor necrosis factor alpha (*TNF-α*) expression at P4 or P14 (Fig S2c-e). Moreover, we did not see changes in the gene expression of *CD3ε* nor *CD19*, which are cell surface markers for T cells and B cells respectively (Fig S4c), suggesting that intestinal lymphocyte reactivity is not altered by DEP/MS exposure. These data again indicate an absence of active inflammation within the small intestine of DEP/MS males in the first two postnatal weeks of life.

In the small intestine, homeostatic macrophage-neuron crosstalk consists of macrophage-derived bone morphogenetic protein 2 (*BMP2*), which activates neurons involved in contractility of the GI tract, in exchange for neuron-derived colony stimulating factor 1 (*CSF1*), essential for macrophage development and survival^37^ (Fig S4d). Furthermore, adrenergic signaling to intestinal macrophages elicits a neuroprotective response from macrophages through an arginase 1 (*ARG1*)-polyamine axis^38^. As such, we determined whether gene expression of these soluble factors is altered in response to DEP/MS exposure. Among macrophage-derived factors, we observed no differences in the male small intestine at P4 or P14 (Fig S4e). However, we observed increased expression of neuron-derived *CSF1* in response to DEP/MS (Fig S4f), mirroring increases in motor neuron genes and *SNAP25* and indicating a coinciding alteration in neuroimmune crosstalk. We did not find any impacts of DEP/MS exposure in intestinal macrophage signaling to smooth muscle cells, where macrophage transient receptor potential vanilloid 4 (*TRPV4*) expression promotes muscular activity through the release of prostaglandin E2 synthesized by prostaglandin E synthase (*PTGES*)^39^ (Fig S4g).

From these results, we infer that secreted factors between macrophages and neurons are not directly mediating DEP/MS perturbations in the male small intestine. However, recent work has shown intestinal macrophages engage in synaptic pruning as a means of neuromodulation—a mechanism previously described only in the central nervous system—specifically at P14 in mouse small intestine^18^. In addition, another group recently found that knocking out macrophage-derived complement component 1q (C1q), a critical mediator of macrophage phagocytosis, leads to robust upregulation of genes associated with neuronal activity and synaptic transmission in the GI tract^40^, consistent with our motor neuron findings at P14. Finally, our group has demonstrated microglia to be deficient in synapse engulfment within the anterior cingulate cortex of the brain, following the same DEP/MS exposure used in this study during the second postnatal week of life in males^12^. Altogether, these data suggested the phagocytic function of intestinal macrophages—rather than their molecular signaling—could be impaired in the male small intestine by DEP/MS exposure, sparing neuronal elements that are normally engulfed at P14.

To investigate this possibility, we repeated the DEP/MS paradigm in a transgenic mouse line in which enhanced green fluorescent protein (eGFP) is expressed under the *CX3CR1* promoter, a common marker for macrophages, efficiently labelling tissue-resident macrophages throughout the small intestine^41^ (**Fig 4a**). This allowed for imaging and 3D reconstructions of intestinal macrophages in P14 *CX3CR1^GFP/WT^* offspring, across all layers of the gut wall (**Fig 4b**). Using this transgenic line, we confirmed that cells positive for *CX3CR1^GFP^* represented the same population of F4/80+ macrophages in the P14 small intestine studied in Figure 2 (Fig S3c-d). We quantified the volume of lysosomal marker CD68 staining within the volume of CX3CR1+ macrophages to determine their phagocytic capacity in males and females exposed to DEP/MS or CON. Simultaneously, we quantified the volume of presynaptic vesicle marker Synapsin1 staining within the lysosome of CX3CR1+ macrophages to measure synaptic engulfment in these same animals. Although we saw no differences in macrophage phagocytic capacity across all layers of the small intestine in response to DEP/MS exposure (**Fig 4c**), we did see significantly less engulfed synaptic material in the muscularis of P14 DEP/MS males compared to CON males (**Fig 4d**). This effect was not present in the submucosal and villi layers of the male small intestine, suggesting that DEP/MS specifically impairs the pruning of myenteric neurons. This is in agreement with the observed upregulation of motor neuron genes (**Fig 3c**). Surprisingly, while we did not see this phenomenon in the muscularis of P14 females, nor in the submucosa, macrophages within the villi of DEP/MS females also engulfed less synaptic material compared to P14 CON females, demonstrating that DEP/MS effects on intestinal macrophages were sex-specific but not exclusive to males (**Fig 4d**). In sum, we saw impaired synaptic engulfment during a critical period of enteric nervous system refinement by intestinal macrophages, in both males and females exposed prenatally to DEP/MS, with our observations in males corresponding spatially and temporally with a postnatal increase in motor neuron transcriptional activity within the small intestine.

**Figure 4.**
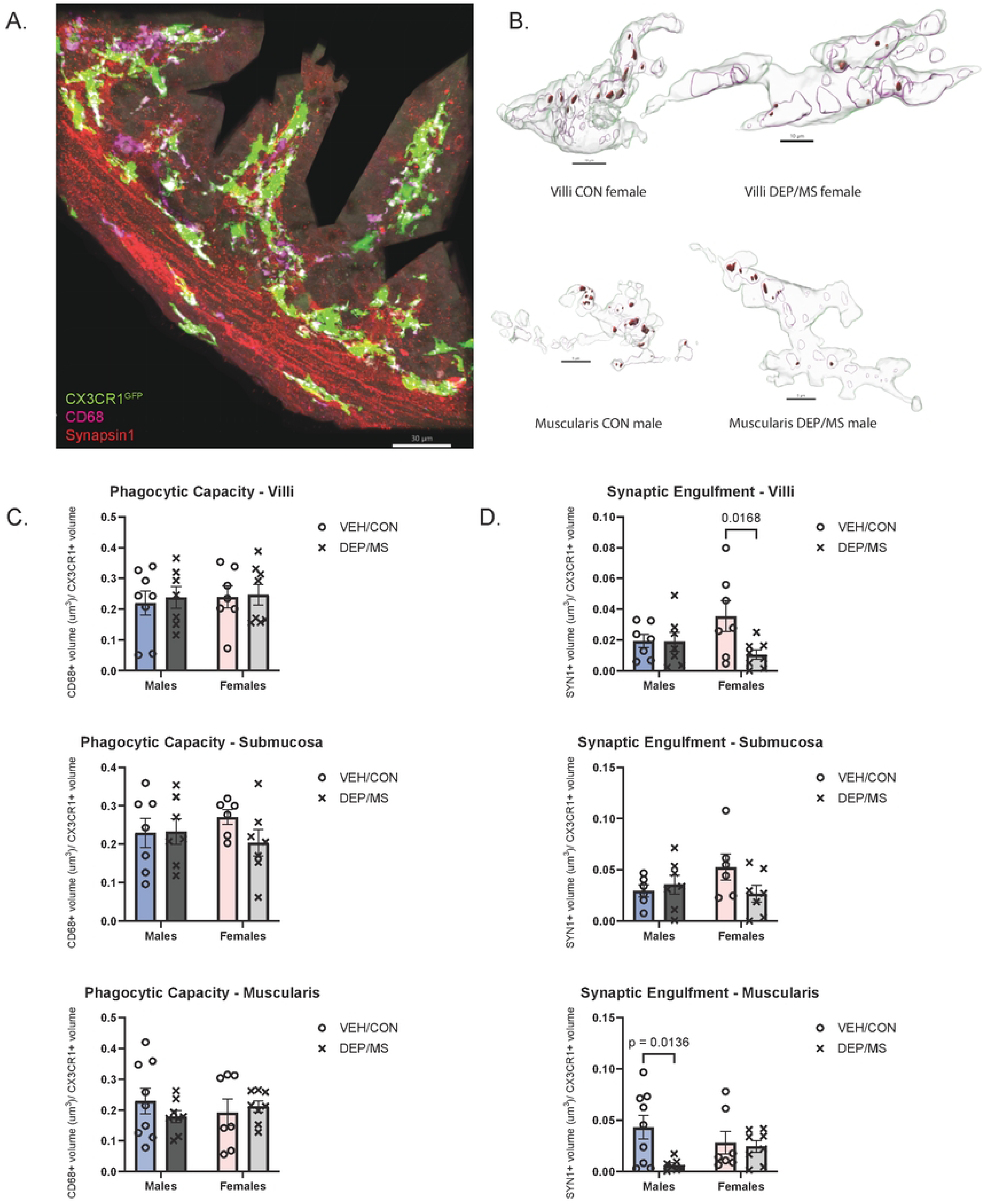
DEP/MS impairs synaptic engulfment by macrophages in the muscularis of male offspring and the villi of female offspring. (A) Representative image of macrophage eGFP expression, lysosomal CD68 staining, and synaptic Synapsin1 staining in P14 CX3CR1GFPJWTtissue. (B) Representative macrophage 3D reconstructions in villi of female offspring and muscularis of male offspring. (C) Quantification of phagocytic capacity (CD68 lysosomal volume *I* eGFP macrophage volume) in P14 offspring in villi (n = 7-8 mice/group, 2-way ANOVA [treatment x sex], treatment: p = 0.7309, sex: p = 0.6923, interaction: p = 0.8769), submucosa (n = 6-7 mice/group, 2-way ANOVA (treatment x sex], treatment: p = 0.3469, sex: p = 0.8584, interaction: p = 0.2943), and muscularis (n = 7-9 mice/group, 2-way ANOVA [treatment x sex], treatment: p = 0.6820, sex: p = 0.9086, interaction: p = 0.3188) (D) Quantification of synaptic engulfment (SYN1 synaptic volume *I* eGFP macrophage volume) in P14 offspring in villi (n = 7-8 mice/group, 2-way ANOVA [treatment x sex], treatment: p = 0.0528, sex: p = 0.5444, interaction: p = 0.0633), submucosa (n = 6-7 mice/group, 2-way ANOVA [treatment x sex], treatment: p = 0.3011, sex: p = 0.4374, interaction: p = 0.0983), and muscularis (n = 7-9 mice/group, 2-way ANOVA [treatment x sex), treatment: p = 0.0327, sex: p = 0.8595, interaction: p = 0.0775)

### Gastrointestinal dysmotility is induced by DEP/MS in P14 males, but not females

Motivated by the convergence of our findings on enteric motor neuron dysregulation following DEP/MS exposure in males, we next assessed the motor function of the GI tract in these offspring. The movement of consumed food and fluids through the GI tract is coordinated by synchronized peristaltic contractions of smooth muscle fibers, and dysfunction of this process is the basis of a large spectrum of digestive issues and GI dysmotility disorders^42^. Moreover, targeted activation of ChAT+ enteric neurons and deletion of macrophage-derived C1q, a mediator of phagocytosis, have both been shown to shorten transit time in the gut^21,40^. Given the prevalence of dysmotility symptoms such as chronic diarrhea or constipation in children with NDDs^8^, and our observations of motor neuron perturbations after DEP/MS exposure, we suspected that DEP/MS exposure may impact transit of luminal contents through the small intestine at the same timepoint. To investigate this, we measured the spatial distribution of FITC-dextran along the GI tract of P14 male and female offspring after oral gavage (**Fig 5a**). Importantly, 45 minutes were taken between gavage and tissue collection of these animals, to constrain fluorescence transit primarily within the small intestine. Indeed, when analyzing both the distribution of fluorescence in these animals and the distribution’s geometric center, we saw that DEP/MS exposure accelerated transit of the fluorophore down the male small intestine (**Fig 5b**). Female offspring did not exhibit any differences in motility following DEP/MS exposure, in agreement with unchanged synaptic engulfment within the P14 muscularis (**Fig 5c**). Taken together, these results demonstrate impaired synaptic engulfment and aberrant motor neuron marker expression within the DEP/MS male small intestine during development, alongside dysregulated gastrointestinal motility.

**Figure 5.**
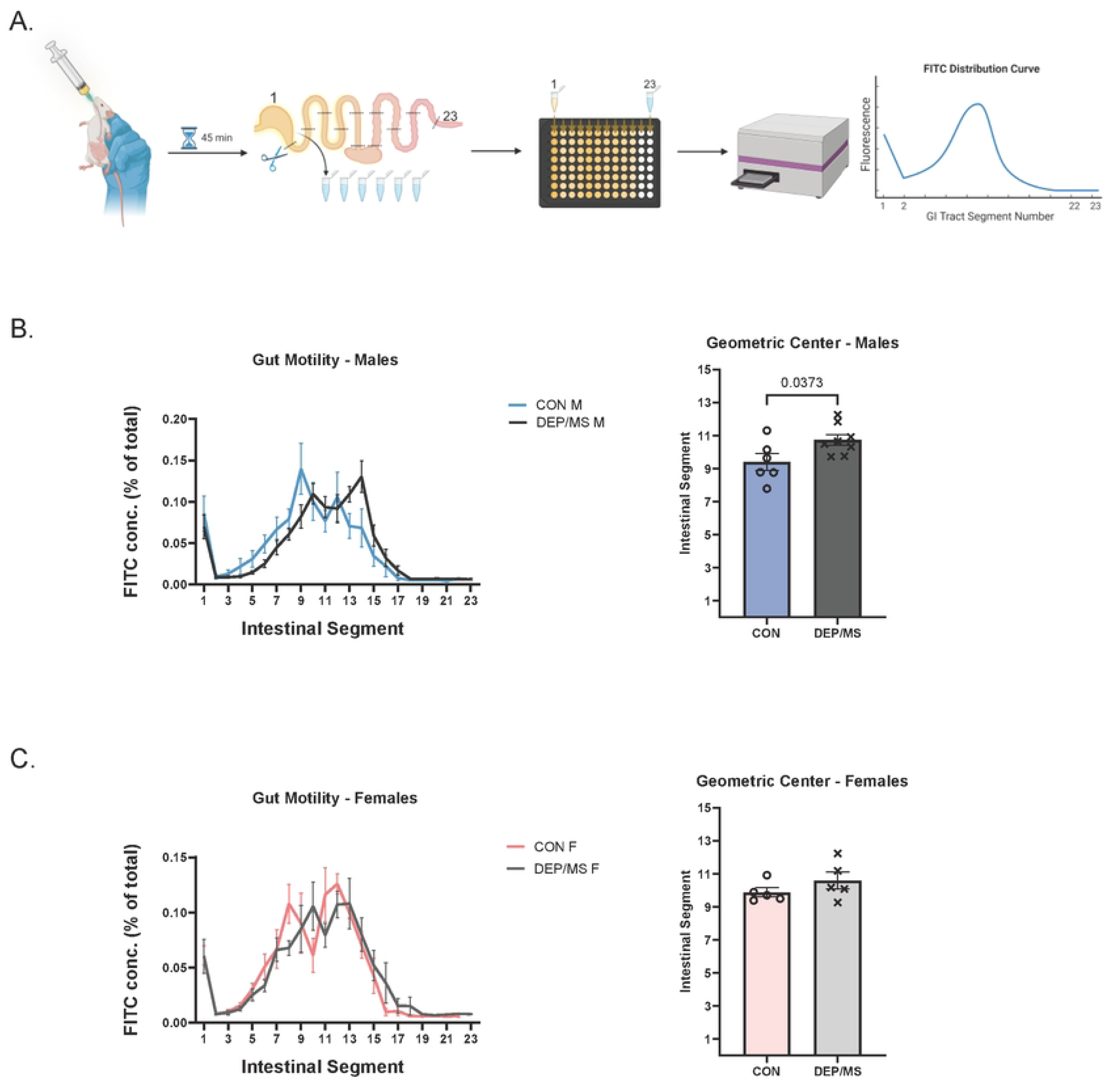
DEP/MS increases gut motility in male offspring. (A) Schematic diagram of gut motility assay. Offspring received FITC-dextran by oral gavage at P14. After 45 minutes, intestine from stomach to rectum was collected, segmented proximally to distally, homogenized along with luminal contents, and measured on a microplate for fluorescent intensity of FITC-dextran distribution along the length of intestine (B) Quantification of FITC-dextran spatial distribution and geometric center in male offspring (n = 6-8 mice/group, unpaired t-test, p = 0.0373) (C) Quantification of FITC-dextran spatial distribution and geometric center in female offspring (n = 5 mice/group, unpaired t-test, p = 0.2687)

## Discussion

Clinical reports of GI comorbidities in children with NDDs encompass a wide range of symptoms including gastritis, bloating, abdominal pain, food intolerances, and dysmotility disorders^27^. While there is little doubt that these abnormalities are mediated in part by the immune system of the gut, mechanistic research has historically focused on inflammatory pathways causing a “leaky gut” post-adolescence^4,43^. This leaves homeostatic roles of intestinal immune cells understudied in the context of development. Here, we report engulfment of synapses within the small intestine—a non-inflammatory function of intestinal macrophages—to be impaired by gestational exposure to DEP/MS, a model representing two major environmental risk factors for NDDs. We found sex-specificity in this response, with male deficits within the muscularis layer of the small intestine corresponding with transcriptional increases in motor neuron activity. Moreover, we show these perturbations to be associated with dysmotility, revealing a novel mechanism by which environmental stressors can directly alter neurodevelopment outside of the brain to impact the health and function of the GI tract.

Developmental insults are known to burden the gut-brain axis^7^, leading to lifelong behavioral deficits and aberrant function of the two organs. While perinatal changes in brain development have been characterized in our previous DEP/MS work^12,25^ and other models of maternal immune activation^4,44,45^, our findings here are some of the first to describe abnormal GI development early in life following gestational challenge. This is especially interesting because numerous studies are finding the gut microbiome to drive neurobehavioral outcomes in NDDs^4,13,46,47^, including our own^6^. However, the origin of microbial differences in models of these disorders remain elusive. During the first weeks of life, maternally-derived microbes colonize the GI tract to form dynamic populations. Given that DEP/MS has no impact on microbiomes within the maternal gut, vagina, or milk^6^, physical perturbations to the developing GI tract, such as dysmotility and villi blunting, may underlie dysbiosis seen in this model, as well as in patients with NDDs^14,47^.

Our findings suggest that DEP/MS specifically impacts muscularis macrophages and their ability to engulf synaptic elements, mirroring impaired synaptic engulfment by microglia observed in this same model^12^. While we observed decreased macrophage marker expression across both muscularis and villi of male DEP/MS offspring, we did not observe reduced synaptic engulfment in the villi. Of note, although the majority of neonatal intestinal macrophages originate from the embryonic yolk sac, villi subpopulations are replaced by bone marrow-derived monocytes while muscularis subpopulations associate with neurons and self-maintain into adulthood^48^. Since bone marrow-derived subpopulations do not carry the fetal programming that yolk sac-derived macrophages carry, impaired synaptic remodeling after DEP/MS might be specific to the muscularis as a result of the longevity of this population. This would coincide with observations in the brain, where microglia are all derived from the embryonic yolk sac and experience limited turnover. We previously observed male-specific impairment of synaptic engulfment by microglia in response to DEP/MS, suggesting a shared mechanism between muscularis macrophages within the small intestine and microglia within the central nervous system^12^. This connection emphasizes a need for further analysis of the ontogeny of long-lived immune cell populations, so that subpopulations with precise functions can be identified early and targeted in therapeutic models of intervention. Furthermore, the impact of early-life stressors outside of the gut-brain axis must be considered, as yolk sac-derived macrophages make their way to all organs throughout the body.

Finally, although DEP/MS induced effects primarily in males, females were not fully spared. Villi macrophages of P14 DEP/MS females engulfed significantly less synaptic material than CON, indicating a similar impairment to that of muscularis macrophages in P14 DEP/MS males. However, the functional consequences of such an abnormality are challenging to dissect due to the great heterogeneity of villi neurons. Afferent connections from these neurons typically lead to higher-order processing in the central nervous system, which lacks clearly defined behavioral outputs for assessment. However, the development of this research would be a boon for our understanding and treatment of sex differences in NDDs, where poor characterization of female-specific phenotypes hinders proper care.

In closing, our results unveil intestinal macrophages to be sensitive to prenatal environmental stressors, dysregulating GI development and function through the enteric nervous system. These players and interactions mirror those found in the brain, implicating yolk sac-derived macrophages as a potential mediator of GI comorbidities in NDDs. As such, our work provides an exciting target for future studies of body-brain axes and therapeutic strategies against GI disorders.

## Methods

### Animals

All procedures were performed in accordance with the guidelines of the National Institutes of Health and Duke University Institutional Animal Care and Use Committee. Wild-type (WT) *C57BL/6J* mice were purchased from Jackson Laboratories (Stock #000664) and *CX3CR1^GFP^* (JAX Stock #008451) mice were obtained from Dr. Ashley Moseman at Duke University. For all experiments, an internal colony of breeding animals was maintained. Animals were group housed in a 12-hour light/dark cycle under standard laboratory conditions.

### Diesel exhaust particle exposure (DEP)

DEP was obtained from Dr. Ian Gilmour at the Environmental Protection Agency and instillations were conducted as previously described^8,12,13^. Briefly, adult females were time mated with the onset of pregnancy indicated by a vaginal plug on embryonic day (E)0. Females were then pair-housed based on embryonic day until E13. Beginning on E2, instillations occurred every 3 days throughout pregnancy, for six total doses. Dams were anesthetized with 2% isoflurane and administered either 50 μg of DEP in 50uL of PBS, 0.05% Tween-20, or vehicle alone (CON) via oropharyngeal aspiration, during which females are suspended by their incisors from a plastic wire and the instillation is administered over the course of 30 s. Females were monitored until they were awake before returning to their home cages.

### Maternal stress exposure (MS)

To induce prenatal maternal stress, we used our previously described^8,12,13^ adaptation of a nest restriction paradigm^26^. On E13, females were singly housed and placed into a cage containing a thin layer of AlphaDri bedding (AlphaDri; Shepherd Specialty Papers) covered by an elevated aluminum mesh platform (0.4 cm × 0.9 cm mesh; McNichols Co., Tampa, FL) and given 2/3 (∼1.9 g) of a cotton nestlet (MS) or placed into a cage with AlphaDri bedding and an entire nestlet (CON). On E18.5 prior to parturition, dams were transferred to a clean cage of AlphaDri bedding with a full nestlet.

### Immunohistochemistry

For all immunohistochemistry experiments, mice were deeply anesthetized via hypothermia or 200 mg/kg tribromoethanol (avertin) before being euthanized via rapid decapitation. 3 cm of the distal small intestine were collected into ice-cold 4% paraformaldehyde (PFA). After 2 hours of PFA fixation, tissue was transferred into 30% sucrose with 0.1% sodium azide for at least 48 hours. Intestines were then embedded in a cryomold filler with O.C.T. (Sakura Finetek) and frozen over dry ice. Using the gut bundling technique, 35 µm transverse sections were made of the intestines using a cryostat (Leica Biosystems) and directly mounted onto positively charged microscope slides. Sections were stored at −20C until staining.

For architectural and macrophage marker staining, WT sections were air-dried at room temperature (RT) for 30 minutes, and then washed in RT PBS three times for 5 minutes each. Following this, tissues were submerged in RT permeabilization buffer (PBS, 1.0% Triton X-100) for 15 minutes. Immediately after, slides were quickly washed in RT PBS and then air-dried at RT briefly, so a hydrophobic barrier could be applied. Sections were then incubated in RT blocking solution (PBS, 5% normal goat serum, 0.5% Tween 20) for 1 hour, before being incubated at 4C with rat anti-F4/80 antibody (1:200 in blocking solution; Abcam ab6640) overnight. Afterwards, tissues were washed in RT PBS three times for 5 minutes each, and then incubated at 4C with Alexa Fluor 594 goat anti-rat IgG (1:200 in blocking solution; Thermo Fisher A-11007) for 2 hours. Slides were washed again in RT PBS three times for 5 minutes each, air-dried at RT, coverslipped with Vectashield Plus mounting medium with 4’,6-diamidino-2-phenylindole (DAPI) (Vector Laboratories H-2000), and stored at 4C until imaging.

For macrophage marker colocalization staining, *CX3CR1^GFP^* sections were stained using the same described methods.

For synaptic engulfment staining, *CX3CR1^GFP^* sections were stained using the same described methods, with substitutions for the appropriate antibodies. During the primary antibody incubation, both rabbit anti-Synapsin1 antibody and rat anti-CD68 antibody (1:100 and 1:250 in blocking solution respectively; Abcam ab64581 and Biolegend 137002 respectively) were used in place of rat anti-F4/80 antibody. During the secondary antibody incubation, both Alexa Fluor 568 goat anti-rabbit IgG and Alexa Fluor 647 goat anti-rat IgG (1:200 in blocking solution for both; Thermo Fisher A-11011 and Thermo Fisher A48265TR respectively) were used in place of Alexa Fluor 594 goat anti-rat IgG.

### Architectural and macrophage marker intensity imaging and analyses

Using a Zeiss microscope with apotome attachment (Axio Imager M1), 20x magnification z-stacks of stained WT small intestines were acquired across 10 focal planes with a 0.5 µm step size. Resultant images were then converted to a maximum intensity projection using Zeiss ZEN software, and then analyzed using QuPath software. To measure the different layers of the small intestine, the polyline tool was used to annotate the transverse cross section based on the visible DAPI morphology, and the length of the annotations averaged across multiple technical replicates from the same animal to give a final value for each animal. Villi were identified by continuous radially inward protrusions of distinct DAPI+ nuclei with no surrounding major artifacts. The muscularis externa was defined radially inwards to include the outermost DAPI+ nuclei of the gut wall and the circumferential band of elongated DAPI+ nuclei, which represent the circular muscularis sublayer. The submucosa was then measured radially outwards, from the base of the villi layer to the start of the muscularis externa layer. Portions of images that did not meet these criteria were omitted from further analysis. To quantify the density of F4/80+ macrophages in each layer of the small intestine, the closed polygon tool was used to first annotate the separates areas of the villi, submucosa, and muscularis. Then, F4/80+ cells were manually counted using the points tool, and the density calculated by dividing the cell count of a layer by the surface area of the layer’s closed polygon annotation. To quantify F4/80 expression levels of these macrophages, somas were each manually selected, and circular ROIs with 5.675 µm radii were generated around each selection. For each ROI, the mean fluorescence intensity for F4/80 was calculated using the measurements feature in QuPath, and then averaged across the F4/80 intensity values for all other ROIs within the same intestinal layer of an image. Finally, macrophage density and F4/80 expression values were averaged across images originating from the same animal to increase the number of technical replicates and provide more accurate representation of the small intestinal tract.

### Macrophage marker colocalization imaging and analysis

Using an Olympus confocal microscope (FLUOVIEW FV3000), 20x magnification 3×3 tile-scans of stained *CX3CR1^GFP^* small intestines were acquired in aross 7 focal planes with a 1.0 µm step size centered within the tissue. Resultant tiles were then stitched together using FV31S-SW Viewer software, and then analyzed using FIJI and ILASTIK software. Z-stack Images were first opened in FIJI to be converted into maximum intensity projections and split into individual channels (eGFP and F4/80). Each channel was saved as a separate TIFF file and transferred to ILASTIK machine learning software for automated identification of cells. ILASTIK was trained using eGFP images from CON and DEP/MS samples to identify eGFP+ cells and create a binary image of these cells. ILASTIK was then trained separately using F4/80 images from CON and DEP/MS samples to identify F4/80+ cells and create a binary image of those cells. For each sample, the binary images from both channels were opened in FIJI, and the number of cells with eGFP signal, F4/80 signal, and overlapping signal was quantified. The final quantification of overlap was made by calculating the percentage of double-positive cells out of the total number of eGFP+ cells.

### Synaptic engulfment imaging and analysis

Using an Olympus confocal microscope (FLUOVIEW FV3000), 60x magnification plus 2.0x optical zoom 2×2 tile-scans of stained *CX3CR1^GFP^* small intestines were acquired across 107 focal planes with a 0.34 µm step size. Resultant tiles were then stitched together using FV31S-SW Viewer software, and then analyzed using FIJI and IMARIS software. Images were first opened in FIJI, so the selection tool could be used to manually separate out the villi, submucosa, and muscularis externa layers with the same methods as used in QuPath. Each layer was then isolated into a separate TIFF file and transferred to IMARIS, for 3D reconstructions of the intestinal macrophages specific to that region of the image. All macrophages present in the field of view were analyzed to maximize technical replicates, with each layer of an animal being represented by at least 3 cells. Surface renderings of intestinal macrophages were created using the CX3CR1^GFP^ signal, and these renderings were used as a mask for the CD68 signal within intestinal macrophages. From this, CD68 surface renderings were then created and used as a mask for the engulfed Synapsin1 signal within macrophage lysosomes. Finally, engulfed Synapsin1 surface renderings were generated, and the volumes of all three surface renderings were extracted using the statistics tool in IMARIS. To calculate phagocytic capacity, the total volume of CD68 was normalized by the total volume of CX3CR1^GFP^. To calculate synaptic engulfment, the total volume of Synapsin1 was normalized by the total volume of CX3CR1^GFP^. This was repeated for each layer of the small intestine for each animal.

### Intestinal permeability

To assess the translocation of inert molecules across the intestinal barrier, P14 mice were food deprived during the first 6 hours of the dark cycle by removing the dam from the home cage and keeping the cage on a 37C heating pad. After this, the body weights of the mice were collected, and then mice were oral gavaged with a solution of 125 mg/mL fluorescein-dextran (FITC-dextran) in PBS at a dosage of 0.6 mg FITC-dextran per gram of body weight. Mice were returned to the heated home cage for 45 minutes, before being anesthetized with 200 mg/kg avertin. Blood was quickly collected from the submandibular vein with a lancet into an untreated tube, allowed to clot for at least 30 minutes, and then spun for 15 minutes at 2000 RCF. Serum was extracted into new tubes, diluted 1:5 in PBS, and then plated in duplicate on an opaque black 96-well microplate. Standards were also plated in duplicate, using FITC-dextran dissolved in 1:5 PBS-gavaged mouse serum:PBS. Standard concentrations of FITC-dextran were 20 µg/mL, 10 µg/mL, 7.5 µg/mL, 5 µg/mL, 4 µg/mL, 3 µg/mL, 2 µg/mL, 1 µg/mL, 0.75 µg/mL, 0.5 µg/mL, 0.25 µg/mL, and 0 µg/mL. Following plating, fluorescence intensity values were read using a CLARIOstar PLUS microplate reader (BMG Labtech), at a 483 nm excitation wavelength and a 535 nm emission wavelength. Raw fluorescence values were then fitted onto a 3^rd^ polynomial regression of the standard curve, and resultant FITC-dextran concentrations were then averaged across duplicates and corrected for dilution with PBS to yield the concentration of circulating FITC-dextran of each P14 mouse.

For P30 mice, the same methods were used with minor adjustments. Mice were weaned at P24 with same-sex siblings and were food deprived prior to the assay by removing all food pellets from the home cage for 2 hours. Mice were oral gavaged with 80 mg/mL of FITC-dextran in PBS at a dosage of 0.6 mg FITC-dextran per gram of body weight. Mice were returned to home cage for 1 hour before rapid submandibular blood collection without avertin.

### qPCR

For all qPCR experiments, mice were deeply anesthetized with 200 mg/kg tribromoethanol (avertin) before being euthanized via rapid decapitation. Small intestines were weighed and collected into a 1.5mL Eppendorf tube, before being flash-frozen and stored at −80C until processing. Tissues were processed directly from −80C storage by being homogenized in 800 µL of 4C Trizol (Thermo Fisher) using a BeadBug 6 homogenizer with 0.6 mm diameter zirconium oxide satellites (Benchmark Scientific). After 15 minutes of resting at RT, 160 µL of chloroform was added to each sample, which were then vortexed for 2 minutes at 2000rpm on an Eppendorf ThermoMixer. Samples were then rested for 3 minutes at RT, before being centrifuged for 15 minutes at 11800 rpm and 4C in an Eppendorf 5415R. From the resulting gradient, the aqueous phase was separated into a new tube, which then received 700 µL of isopropanol to precipitate RNA. Samples were again vortexed for 2 minutes at 2000 rpm, rested for 10 minutes at RT, and then centrifuged at the same settings to form pellets. Pellets were rinsed twice in 4C 75% ethanol and then resuspended in 80-120 µL of nuclease-free water. RNA was frozen at −80C until cDNA synthesis using the QuantiTect Reverse Transcription Kit (Qiagen 205314). Briefly, RNA (2000 ng per 24 µl nuclease-free water) from each sample was pre-treated with gDNase at 42C for 2 minutes to remove genomic DNA contamination. Next, master mix containing both primer-mix and reverse transcriptase was added to each sample and all samples were heated to 42C for 30 minutes and then 95C for 3 minutes in the thermocycler. qPCR was run on a Mastercycler ep realplex system (Eppendorf) using the QuantiNova SYBR Green PCR Kit (Qiagen 208057). All PCR primers were designed in the lab or from the Harvard PrimerBank, and the oligonucleotides were purchased from Integrated DNA technologies. Relative gene expression was calculated using the 2^-ΔΔCT^ method, relative to the house-keeping gene *18S* and the average ΔCT of the P4 CON male samples. Samples were removed before unblinding if they failed to amplify or if there was a secondary peak in the melting temperature plot indicating contamination.

Primer sequences for all genes are as follows from 5’ to 3’:

***18S****, F: GAA TAA TGG AAT AGG ACC GC, R: CTT TCG CTC TGG TCC GTC TT;*

***SNAP25****, F: CAA CTG GAA CGC ATT GAG GAA, R: GGC CAC TAC TCC ATC CTG ATT AT;*

***ChAT****, F: CCA TTG TGA AGC GGT TTG GG, R: GCC AGG CGG TTG TTT AGA TAC A;*

***nNOS****, F: CTG GTG AAG GAA CGG GTC AG, R: CCG ATC ATT GAC GGC GAG AAT;*

***VGLUT2****, F: TGG AAA ATC CCT CGG ACA GAT, R: CAT AGC GGA GCC TTC TTC TCA;*

***CGRP****, F: GGA CTT GGA GAC AAA CCA CCA, R: GAG AGC AAC CAG AGA GGA ACT ACA;*

***TPH2****, F: GGT TGT CCT TGG ATT CTG CTG, R: GCC TGG ATT CGA TAT GAA GCA T;*

***TH****, F: GTC TCA GAG CAG GAT ACC AAG C, R: CTC TCC TCG AAT ACC ACA GCC;*

***DBH****, F: GAG GCG GCT TCC ATG TAC G, R: TCC AGG GGG ATG TGG TAG G;*

***MuSK****, F: TGA GAA CTG CCC CTT GGA ACT, R: GGG TCT ATC AGC AGG CAG CTT;*

***CD3e****, F: ATG CGG TGG AAC ACT TTC TGG, R: GCA CGT CAA CTC TAC ACT GGT;*

***CD19****, F: GGA GGC AAT GTT GTG CTG C, R: ACA ATC ACT AGC AAG ATG CCC;*

***BMP2****, F: GGG ACC CGC TGT CTT CTA GT, R: TCA ACT CAA ATT CGC TGA GGA C;*

***Arg1****, F: CTC CAA GCC AAA GTC CTT AGA G, R: AGG AGC TGT CAT TAG GGA CAT C;*

***CSF1****, F: ATG AGC AGG AGT ATT GCC AAG G, R: TCC ATT CCC AAT CAT GTG GCT A;*

***TRPV4****, F: CCT GCT GGT CAC CTA CAT CA, R: CTC AGG AAC ACA GGG AAG GA;*

***PTGES****, F: GGA TGC GCT GAA ACG TGG A, R: CAG GAA TGA GTA CAC GAA GCC*

### Intestinal motility

To measure the spatial migration of an inert molecule along the GI tract, P14 mice were food deprived during the first 6 hours of the dark cycle by removing the dam from the home cage and keeping the cage on a 37C heating pad. After this, the body weights of the mice were collected, and then mice were oral gavaged with a solution of 125 mg/mL FITC-dextran in PBS at a dosage of 0.6 mg FITC-dextran per gram of body weight. Mice were returned to the heated home cage for 45 minutes, before being anesthetized with 200 mg/kg avertin and subsequently euthanized via rapid decapitation. From the stomach to the rectum, GI tracts were carefully extracted onto a clean, dry petri dish, where the lengths of the small intestine and overall GI tract were measured with a ruler. Then, entire GI tracts were cut with a razor blade into equal length pieces, numbered up proximal to distal to 23, which were individually submerged into clean tubes filled with 1 mL of PBS. Intestinal segments were then homogenized along with their luminal contents in these tubes, using a sonicator, and then centrifuged at 1000 rcf for 10 minutes. Supernatants were then diluted 1:99 in PBS and plated in duplicate on an opaque black 96-well microplate, along with standards made from FITC-dextran dissolved in PBS. Standard concentrations were 30 µg/mL, 15 µg/mL, 10 µg/mL, 9 µg/mL, 8 µg/mL, 7 µg/mL, 6 µg/mL, 5 µg/mL, 4 µg/mL, 3 µg/mL, 2 µg/mL, and 0 µg/mL. Following plating, fluorescence intensity values were read using a CLARIOstar PLUS microplate reader, at a 483 nm excitation wavelength and a 535 nm emission wavelength. Raw fluorescence values were then fitted onto a 3^rd^ polynomial regression of the standard curve, and resultant FITC-dextran concentrations were then averaged across duplicates and corrected for dilution with PBS to yield the raw amount of FITC-dextran within each intestinal segment. Finally, these values were normalized by the total amount of FITC-dextran across all the intestinal segments of the same mouse, giving the percentage of FITC-dextran contained within each intestinal segment relative to the entire GI tract. Geometric centers of this spatial distribution were calculated for each individual mouse by multiplying these percentages with their respective segment number, and then summing together all the resultant values within that mouse. Σ(% of total fluorescent signal per segment × segment number).

### Statistical analysis

Statistical tests were performed using GraphPad Prism 10. Due to our sample sizes and the need to analyze eight different groups (CON vs. DEP/MS, P4 vs. P14, Male vs. Female), data are presented and analyzed segregated by sex to avoid type II (false negative) results when all eight groups are present^49^. Unless otherwise noted, two-way ANOVAs were performed on data sets with two independent variables (eg sex x treatment, age x treatment), along with Sidak’s post-hoc analysis. Unpaired t-tests were performed on data sets with one independent variable. Data are expressed as mean ± SEM and statistical significance was set at *p* < 0.05.

## Acknowledgements

We would like to thank all the members of the Bilbo lab for many valuable years of discussions and feedback. We also would like to thank the animal care staff at Duke University for making this work possible. We thank Dr. Ian Gilmour of the Environmental Protection Agency for the DEP. We thank the Duke Light Microscopy Core Facility for access to software for image analysis. BioRender was used to generate main figure schematics.

